# Absence of SARS-CoV-2 in Wildlife of Northeastern Minnesota and Isle Royale National Park

**DOI:** 10.1101/2023.07.14.549077

**Authors:** David Castañeda, Edmund J. Isaac, Todd Kautz, Mark C. Romanski, Seth A. Moore, Matthew T. Aliota

## Abstract

We investigated the presence of SARS-CoV-2 in free-ranging wildlife populations in Northeastern Minnesota on the Grand Portage Indian Reservation and Isle Royale National Park. 120 nasal samples were collected from white-tailed deer, moose, gray wolves, and black bears monitored for conservation efforts during 2022-2023. Samples were tested for viral RNA by RT-qPCR using the CDC N1/N2 primer set. Our data indicate that no wildlife samples were positive for SARS-CoV-2 RNA. Continued surveillance is therefore crucial to better understand the changing landscape of zoonotic SARS-CoV-2 in the Upper Midwest.

## Main Text

Severe acute respiratory syndrome coronavirus 2 (SARS-CoV-2)—the etiologic agent of Coronavirus Disease-19 (COVID-19)—is believed to have emerged from an animal source and has subsequently been identified as zoonotic (Lu et al., 2020; Zhou et al., 2020). However, the exact species involved in the virus’s origin and its intermediate host have yet to be identified (Alwine et al., 2023; Zhao et al., 2020). Still, SARS-CoV-2 is capable of infecting a wide range of mammalian species. Multiple reports have documented evidence of infection in cats (*Felis catus*), dogs (*Canis lupus familiaris*), hamsters (*Mesocricetus auratus*), ferrets (*Mustela putorius furo*), farmed minks (*Neovison vison*), raccoon dogs (*Nyctereutes porcyonoides*), tigers (*Panthera tigris*), snow leopards (*Panthera uncia*), pumas (*Puma concolor*), lions (*Panthera leo*), rhesus macaques (*Macaca mulatta*), gorillas (*Gorilla gorilla*), rabbits (*Oryctolagus cuniculus*), and fruit bats (*Rousettus aegyptiacus*) (Freuling et al., 2020; Gortázar et al., 2021; Hobbs & Reid, 2021; Jo et al., 2021; Muñoz-Fontela et al., 2020; Sharun et al., 2021; Shi et al., 2020), among others.

Recently it has been shown that white-tailed deer (*Odocoileus virginianus*), are highly susceptible to infection with SARS-CoV-2, are exposed to a range of viral diversity from humans, and are capable of sustaining transmission in nature (Feng et al., 2023; Kuchipudi et al., 2022; Willgert et al., 2022). The detection of SARS-CoV-2 in free-ranging white-tailed deer, therefore, naturally raises the question whether less accessible species are also being infected through viral spillover from humans or by a bridge species like white-tailed deer. In wildlife settings, deer often encounter other related species such as moose (*Alces alces*) and gray wolves (*Canis lupus*), especially in northern Minnesota, where they share sympatric habitats. Studies have reported a dynamic predator-prey relationship between moose, deer, and wolves, which can lead to spillover events involving parasitic pathogens (Oliveira-Santos et al., 2021). Therefore, it is plausible to consider them as a potential source for SARS-CoV-2 transmission to moose, gray wolves, and other species within shared geographic foci. Furthermore, it has previously been shown that residues in the ACE2 receptor associated with SARS-CoV-2 susceptibility are highly conserved among cervids, with moose and deer having identical residues that classify them as high-risk species (Lopes, 2023; Damas et al. in 2020). All of which argue for broader surveillance efforts.

Moose in Minnesota are found in the northeastern part of the state where the population has declined from about 10,000 individuals in 2006 to about 3290 currently (2023) (Giudice, 2023). Moose are a critical food source and an integral part of spiritual and cultural practices for the Grand Portage Band of Lake Superior Chippewa (GPBLSC), which is a federally recognized Indian tribe in extreme northeastern Minnesota at the urban/wildlife interface. The GPBLSC proudly exercises its rights to food sovereignty through subsistence hunting and fishing, and moose and deer are the primary terrestrial subsistence food used by the GPBLSC. As a result, the GPBLSC is heavily engaged in conservation efforts to maintain the moose population in the state. Subsistence hunting could create a scenario for SARS-CoV-2 spillover into wildlife and/or spillback into the indigenous community. This is particularly concerning because indigenous communities have experienced substantially greater rates of COVID-19 mortality compared with other racial and ethnic groups (Leggat-Barr et al., 2021). To better understand zoonotic SARS-CoV-2 in Minnesota, we examined the prevalence of SARS-CoV-2 RNA in four wildlife species that are important to the GPBLSC using reverse transcription quantitative polymerase-chain reaction (RT-qPCR). We sampled 27 white-tailed deer, 78 moose, 8 gray wolves (*Canis lupus*), and 7 black bears (*Ursus americanus*) during conservation efforts in 2022 and 2023 across three different locations: Grand Portage and Isabella in northeastern Minnesota, and Isle Royale National Park, Michigan.

Nasal swabs were collected using the DNA/RNA Shield SafeCollect Swab Collection Kit from Zymo Research (R1007e). Nasal specimens were collected by inserting a dry, sterile swab 8-12 cm into each nostril and gently making five passes around the interior of the nostril. If the nasal cavity was inaccessible, throat swabs were taken by inserting the swab as far back into the throat as possible. Samples were extracted using the Quick-RNA Viral Kit (Zymo Research). The extraction method followed manufacturer-recommended protocols with the notable exceptions of using 100□μL of starting material and eluting with 65□μL of appropriate elution material as indicated by manufacturer protocols. RT-qPCR reactions were set up in a 96-well Barcoded plate (Thermo Fisher Scientific) for either the N1 or N2 primers and probes with CDC-recommended sequences. Samples with Ct □ < □ 40 in both N1 and N2 reactions were determined as positive for SARS-CoV-2. In contrast, samples with Ct □ < □ 40 in only N1 or N2 targets were determined as inconclusive results, and samples without amplification in both N1 and N2 targets were deemed negative for SARS-CoV-2. We found no detectable SARS-CoV-2 RNA within moose, deer, wolf, or bear samples. Positive control samples on each plate amplified as expected. We did detect SARS-CoV-2 RNA in 1/27 deer samples with high N1 and N2 CTs, but this sample was also positive for human Rnase P, thus it is not possible to determine if this signal is derived from a deer or human infection.

In sum, our study did not find evidence for SARS-CoV-2 infection in white-tailed deer, moose, gray wolves, or black bears. However, this does not mean that they are not susceptible. Indeed, numerous reports have documented spillover events from humans to deer populations in North America (Chandler et al., 2021; Feng et al., 2023; Hale et al., 2022; Hancock et al., 2022; Kuchipudi et al., 2022; Marques et al., 2022; Palermo et al., 2021; Pickering et al., 2022; Roundy et al., 2022; Vandegrift et al., 2022). For example, a study encompassing Michigan, Illinois, New York, and Pennsylvania found that 40% of free-ranging deer tested positive for SARS-CoV-2 antibodies (Chandler et al., 2021). In another study conducted in Ohio, approximately one-third of the deer population tested positive for the virus using RT-qPCR, and sequencing provided evidence for potential deer-to-deer transmission (Hale et al., 2022). Genomic surveillance in Ontario, Canada also supported deer-to-deer transmission and also provided strong evidence suggesting deer-to-human transmission (Pickering et al., 2022).

It is also important to note that other studies suggest that infections in wildlife appear to be highly focal and sporadic. For instance, in Alabama, approximately 100 captive deer were sampled, and while humans in close contact with the deer tested positive for SARS-CoV-2— none of the deer samples were positive (Barua et al., 2022). Similarly, a Canadian study expanded wildlife surveillance to 17 different species, including deer, black bears, and wolves, but found no evidence of SARS-CoV-2 RNA or antibodies in any of the animals (Greenhorn et al., 2022)—similar results were recently observed in Vermont as well (Despres et al., 2023). In Europe, studies conducted in the UK, Germany, and Austria analyzing different deer species endemic to the region, showed no evidence of exposure or susceptibility to SARS-CoV-2 (Holding et al., 2022; Moreira-Soto et al., 2022).

Finally, there are several important factors that could have influenced the results reported herein. Deer are social animals and females travel in herds and have frequent contact with peri-urban and urban human populations. In contrast, moose are usually solitary and do not travel in herds. As a result, there is less opportunity for moose to share pathogens with humans and/or other animal species. Similarly, Grand Portage, where most of our samples originated, has a very low human population density and is relatively remote and therefore not connected to adjacent urban areas. Only a few samples were collected during the beginning of the hunting season in 2022, while the remaining samples were taken during winter, when there is likely less human contact with wildlife populations in those areas. Reduced human contact with animals decreases the chances of spillover to these populations, which might have been a main factor in the absence of cases detected in comparison to other studies. Our goal was to expand surveillance efforts and explore other related species that have contact with deer, nonetheless, increasing the number of deer samples compared to the other species, might have increased our chances of finding positive results.

While it remains unclear whether moose, black bear, or gray wolf are susceptible to SARS-CoV-2, our findings suggest that transmission of SARS-CoV-2 in Northeastern Minnesota and Isle Royale National Park wildlife is not common. Still, these data argue for continued studies (experimental and epidemiologic) to investigate the potential susceptibility of these and other wildlife species to SARS-CoV-2. Monitoring wildlife populations for the presence of SARS-CoV-2 can help identify potential reservoir hosts and assess putative pathways for viral spillover into wildlife. To determine the risk of zoonotic SARS-CoV-2 transmission and the establishment of sylvatic SARS-CoV-2, where the constraints on evolution may be different than what occurs in humans, surveillance of diverse animal species will be paramount.

## Acknowledgments

We thank Grand Portage Band of Lake Superior fish and wildlife technicians Roger (Poe) Deschampe and Frank Manthy for assistance in capturing animals. We acknowledge Heliwild Helicopter Wildlife Services for supplying helicopters for this work. Funding for this project came from start-up funds from the University of Minnesota Department of Veterinary and Biomedical Sciences to MTA.

---

**Table 1.** Samples analyzed from different species during 2022 and 2023.

| **Location** | **Species** | **No. of samples in**<br>**2022** | **No. of Samples in**<br>**2023** |
| --- | --- | --- | --- |
| **Grand**<br>**Portage** | White-tailed<br>deer | 14 | 13 |
|  | Moose | 19 | 23 |
|  | Bear | 7 | - |
|  | Wolf | 3 | - |
| **Isabella** | Moose | 10 | - |
| **Isle Royale** | Moose | 19 | 7 |
|  | Wolf | 5 | - |
|  | **Total** | **77** | **43** |

## References

Alwine, J. C., Casadevall, A., Enquist, L. W., Goodrum, F. D., & Imperiale, M. J. (2023). A Critical Analysis of the Evidence for the SARS-CoV-2 Origin Hypotheses. Journal of Virology, 97(4), e00365–23. https://doi.org/10.1128/jvi.00365-23

Barua, S., Newbolt, C. H., Ditchkoff, S. S., Johnson, C., Zohdy, S., Smith, R., & Wang, C. (2022). Absence of SARS-CoV-2 in a captive white-tailed deer population in Alabama, USA. Emerging Microbes & Infections, 11(1), 1707–1710. https://doi.org/10.1080/22221751.2022.2090282

Caserta, L. C., Martins, M., Butt, S. L., Hollingshead, N. A., Covaleda, L. M., Ahmed, S., Everts, M. R. R., Schuler, K. L., & Diel, D. G. (2023). White-tailed deer (Odocoileus virginianus) may serve as a wildlife reservoir for nearly extinct SARS-CoV-2 variants of concern. Proceedings of the National Academy of Sciences, 120(6), e2215067120. https://doi.org/10.1073/pnas.2215067120

Chandler, J. C., Bevins, S. N., Ellis, J. W., Linder, T. J., Tell, R. M., Jenkins-Moore, M., Root, J. J., Lenoch, J. B., Robbe-Austerman, S., DeLiberto, T. J., Gidlewski, T., Kim Torchetti, M., & Shriner, S. A. (2021). SARS-CoV-2 exposure in wild white-tailed deer (Odocoileus virginianus). Proceedings of the National Academy of Sciences, 118(47), e2114828118. https://doi.org/10.1073/pnas.2114828118

Damas, J., Hughes, G. M., Keough, K. C., Painter, C. A., Persky, N. S., Corbo, M., Hiller, M., Koepfli, K.-P., Pfenning, A. R., Zhao, H., Genereux, D. P., Swofford, R., Pollard, K. S., Ryder, O. A., Nweeia, M. T., Lindblad-Toh, K., Teeling, E. C., Karlsson, E. K., & Lewin, H. A. (2020). Broad host range of SARS-CoV-2 predicted by comparative and structural analysis of ACE2 in vertebrates. Proceedings of the National Academy of Sciences, 117(36), 22311–22322. https://doi.org/10.1073/pnas.2010146117

Despres, H. W., Mills, M. G., Schmidt, M. M., Gov, J., Perez, Y., Jindrich, M., Crawford, A. M. L., Kohl, W. T., Rosenblatt, E., Kubinski, H. C., Simmons, B. C., Nippes, M. C., Goldenberg, A. J., Murtha, K. E., Nicoloro, S., Harris, M. J., Feeley, A. C., Gelinas, T. K., Cronin, M. K., … Bruce, E. A. (2023). Surveillance of Vermont wildlife in 2021-2022 reveals no detected SARS-CoV-2 viral RNA [Preprint]. Microbiology. https://doi.org/10.1101/2023.04.25.538264

Feng, A., Bevins, S. N., Chandler, J., DeLiberto, T. J., Ghai, R., Lantz, K., Lenoch, J., Retchless, A., Shriner, S. A., Tang, C. Y., Tong, S. S., Torchetti, M., Uehara, A., & Wan, X. (2023). Transmission of SARS-CoV-2 in free-ranging white-tailed deer in the United States. Nature Communications, 14(1). https://doi.org/10.1038/s41467-023-39782-x

Freuling, C. M., Breithaupt, A., Müller, T., Sehl, J., Balkema-Buschmann, A., Rissmann, M., Klein, A., Wylezich, C., Höper, D., Wernike, K., Aebischer, A., Hoffmann, D., Friedrichs, V., Dorhoi, A., Groschup, M. H., Beer, M., & Mettenleiter, T. C. (2020). Susceptibility of Raccoon Dogs for Experimental SARS-CoV-2 Infection. Emerging Infectious Diseases, 26(12), 2982–2985. https://doi.org/10.3201/eid2612.203733

Giudice, J. H., MNDNR Wildlife Biometrics Group. (2023). 2023 AERIAL MOOSE SURVEY. https://files.dnr.state.mn.us/wildlife/moose/moosesurvey.pdf

Gortázar, C., Barroso-Arévalo, S., Ferreras-Colino, E., Isla, J., de la Fuente, G., Rivera, B., Domínguez, L., de la Fuente, J., & Sánchez-Vizcaíno, J. M. (2021). Natural SARS-CoV-2 Infection in Kept Ferrets, Spain. Emerging Infectious Diseases, 27(7), 1994–1996. https://doi.org/10.3201/eid2707.210096

Greenhorn, J., Kotwa, J., Bowman, J., Bruce, L., Buchanan, T., Buck, P., Davy, C., Dibernardo, A., Flockhart, L., Gagnier, M., Hou, A., Jardine, C., Lair, S., Lindsay, L., Massé, A., Muchaal, P., Nituch, L., Sotto, A., Stevens, B., … Mubareka, S. (2022). SARS-CoV-2 wildlife surveillance in Ontario and Québec. Canada Communicable Disease Report, 48(6), 243–251. https://doi.org/10.14745/ccdr.v48i06a02

Hale, V. L., Dennis, P. M., McBride, D. S., Nolting, J. M., Madden, C., Huey, D., Ehrlich, M., Grieser, J., Winston, J., Lombardi, D., Gibson, S., Saif, L., Killian, M. L., Lantz, K., Tell, R. M., Torchetti, M., Robbe-Austerman, S., Nelson, M. I., Faith, S. A., & Bowman, A. S. (2022). SARS-CoV-2 infection in free-ranging white-tailed deer. Nature, 602(7897), 481–486. https://doi.org/10.1038/s41586-021-04353-x

Hancock, T. J., Hickman, P., Kazerooni, N., Kennedy, M., Kania, S. A., Dennis, M., Szafranski, N., Gerhold, R., Su, C., Masi, T., Smith, S., & Sparer, T. E. (2022). Possible Cross-Reactivity of Feline and White-Tailed Deer Antibodies against the SARS-CoV-2 Receptor Binding Domain. Journal of Virology, 96(8), e00250–22. https://doi.org/10.1128/jvi.00250-22

Hobbs, E. C., & Reid, T. J. (2021). Animals and SARS □ CoV □ 2: Species susceptibility and viral transmission in experimental and natural conditions, and the potential implications for community transmission. Transboundary and Emerging Diseases, 68(4), 1850–1867. https://doi.org/10.1111/tbed.13885

Holding, M., Otter, A. D., Dowall, S., Takumi, K., Hicks, B., Coleman, T., Hemingway, G., Royds, M., Findlay Wilson, S., Curran French, M., Vipond, R., Sprong, H., & Hewson, R. (2022). Screening of wild deer populations for exposure to SARS □ CoV □ 2 in the United Kingdom, 2020–2021. Transboundary and Emerging Diseases, 69(5). https://doi.org/10.1111/tbed.14534

Jo, W. K., Oliveira□Filho, E. F., Rasche, A., Greenwood, A. D., Osterrieder, K., & Drexler, J. F. (2021). Potential zoonotic sources of SARS □ CoV □ 2 infections. Transboundary and Emerging Diseases, 68(4), 1824–1834. https://doi.org/10.1111/tbed.13872

Kuchipudi, S. V., Surendran-Nair, M., Ruden, R. M., Yon, M., Nissly, R. H., Vandegrift, K. J., Nelli, R. K., Li, L., Jayarao, B. M., Maranas, C. D., Levine, N., Willgert, K., Conlan, A. J. K., Olsen, R. J., Davis, J. J., Musser, J. M., Hudson, P. J., & Kapur, V. (2022). Multiple spillovers from humans and onward transmission of SARS-CoV-2 in white-tailed deer. Proceedings of the National Academy of Sciences, 119(6), e2121644119. https://doi.org/10.1073/pnas.2121644119

Lan, J., Ge, J., Yu, J., Shan, S., Zhou, H., Fan, S., Zhang, Q., Shi, X., Wang, Q., Zhang, L., & Wang, X. (2020). Structure of the SARS-CoV-2 spike receptor-binding domain bound to the ACE2 receptor. Nature, 581(7807), 215–220. https://doi.org/10.1038/s41586-020-2180-5

Leggat-Barr, K., Uchikoshi, F., & Goldman, N. (2021). COVID-19 risk factors and mortality among Native Americans. Demographic Research, 45, 1185–1218. https://doi.org/10.4054/DemRes.2021.45.39

Li, W., Moore, M. J., Vasilieva, N., Sui, J., Wong, S. K., Berne, M. A., Somasundaran, M., Sullivan, J. L., Luzuriaga, K., Greenough, T. C., Choe, H., & Farzan, M. (2003). Angiotensin-converting enzyme 2 is a functional receptor for the SARS coronavirus. Nature, 426(6965), 450–454. https://doi.org/10.1038/nature02145

Lopes, L. R. (2023). Cervids ACE2 Residues that Bind the Spike Protein can Provide Susceptibility to SARS-CoV-2. EcoHealth, 20(1), 9–17. https://doi.org/10.1007/s10393-023-01632-z

Lu, R., Zhao, X., Li, J., Niu, P., Yang, B., Wu, H., Wang, W., Song, H., Huang, B., Zhu, N., Bi, Y., Ma, X., Zhan, F., Wang, L., Hu, T., Zhou, H., Hu, Z., Zhou, W., Zhao, L., … Tan, W. (2020). Genomic characterisation and epidemiology of 2019 novel coronavirus: Implications for virus origins and receptor binding. The Lancet, 395(10224), 565–574. https://doi.org/10.1016/S0140-6736(20)30251-8

Marques, A. D., Sherrill-Mix, S., Everett, J. K., Adhikari, H., Reddy, S., Ellis, J. C., Zeliff, H., Greening, S. S., Cannuscio, C. C., Strelau, K. M., Collman, R. G., Kelly, B. J., Rodino, K. G., Bushman, F. D., Gagne, R. B., & Anis, E. (2022). Multiple Introductions of SARS-CoV-2 Alpha and Delta Variants into White-Tailed Deer in Pennsylvania. MBio, 13(5), e02101–22. https://doi.org/10.1128/mbio.02101-22

Minnesota Department of Health. (2023). COVID-19 Vaccine Data COVID-19 Situation Update. https://www.health.state.mn.us/diseases/coronavirus/stats/vaccine.html

Minnesota Department of Natural Resources. (2018). Minnesota White-tailed deer Management Plan. https://files.dnr.state.mn.us/wildlife/deer/plan/deerplan.pdf

Moreira-Soto, A., Walzer, C., Czirják, G. Á., Richter, M. H., Marino, S. F., Posautz, A., De Yebra Rodo, P., McEwen, G. K., Drexler, J. F., & Greenwood, A. D. (2022). Serological Evidence That SARS-CoV-2 Has Not Emerged in Deer in Germany or Austria during the COVID-19 Pandemic. Microorganisms, 10(4), 748. https://doi.org/10.3390/microorganisms10040748

Muñoz-Fontela, C., Dowling, W. E., Funnell, S. G. P., Gsell, P.-S., Riveros-Balta, A. X., Albrecht, R. A., Andersen, H., Baric, R. S., Carroll, M. W., Cavaleri, M., Qin, C., Crozier, I., Dallmeier, K., de Waal, L., de Wit, E., Delang, L., Dohm, E., Duprex, W. P., Falzarano, D., … Barouch, D. H. (2020). Animal models for COVID-19. Nature, 586(7830), 509–515. https://doi.org/10.1038/s41586-020-2787-6

Oliveira-Santos, L. G. R., Moore, S. A., Severud, W. J., Forester, J. D., Isaac, E. J., Chenaux-Ibrahim, Y., Garwood, T., Escobar, L. E., & Wolf, T. M. (2021). Spatial compartmentalization: A nonlethal predator mechanism to reduce parasite transmission between prey species. Science Advances, 7(52), eabj5944. https://doi.org/10.1126/sciadv.abj5944

Palermo, P. M., Orbegozo, J., Watts, D. M., & Morrill, J. C. (2021). SARS-CoV-2 Neutralizing Antibodies in White-Tailed Deer from Texas. Vector-Borne and Zoonotic Diseases, vbz.2021.0094. https://doi.org/10.1089/vbz.2021.0094

Pappas, G., Vokou, D., Sainis, I., & Halley, J. M. (2022). SARS-CoV-2 as a Zooanthroponotic Infection: Spillbacks, Secondary Spillovers, and Their Importance. Microorganisms, 10(11), 2166. https://doi.org/10.3390/microorganisms10112166

Pickering, B., Lung, O., Maguire, F., Kruczkiewicz, P., Kotwa, J. D., Buchanan, T., Gagnier, M., Guthrie, J. L., Jardine, C. M., Marchand-Austin, A., Massé, A., McClinchey, H., Nirmalarajah, K., Aftanas, P., Blais-Savoie, J., Chee, H.-Y., Chien, E., Yim, W., Banete, A., … Bowman, J. (2022). Divergent SARS-CoV-2 variant emerges in white-tailed deer with deer-to-human transmission. Nature Microbiology, 7(12), 2011–2024. https://doi.org/10.1038/s41564-022-01268-9

Roundy, C. M., Nunez, C. M., Thomas, L. F., Auckland, L. D., Tang, W., Richison, J. J., Green, B. R., Hilton, C. D., Cherry, M. J., Pauvolid-Corrêa, A., Hamer, G. L., Cook, W. E., & Hamer, S. A. (2022). High Seroprevalence of SARS-CoV-2 in White-Tailed Deer (Odocoileus virginianus) at One of Three Captive Cervid Facilities in Texas. Microbiology Spectrum, 10(2), e00576–22. https://doi.org/10.1128/spectrum.00576-22

Sharun, K., Dhama, K., Pawde, A. M., Gortázar, C., Tiwari, R., Bonilla-Aldana, D. K., Rodriguez-Morales, A. J., de la Fuente, J., Michalak, I., & Attia, Y. A. (2021). SARS-CoV-2 in animals: Potential for unknown reservoir hosts and public health implications. Veterinary Quarterly, 41(1), 181–201. https://doi.org/10.1080/01652176.2021.1921311

Shi, J., Wen, Z., Zhong, G., Yang, H., Wang, C., Huang, B., Liu, R., He, X., Shuai, L., Sun, Z., Zhao, Y., Liu, P., Liang, L., Cui, P., Wang, J., Zhang, X., Guan, Y., Tan, W., Wu, G., … Bu, Z. (2020). Susceptibility of ferrets, cats, dogs, and other domesticated animals to SARS–coronavirus 2. Science, 368(6494), 1016–1020. https://doi.org/10.1126/science.abb7015

Sparrer, M. N., Hodges, N. F., Sherman, T., VandeWoude, S., Bosco-Lauth, A. M., & Mayo, C. E. (2023). Role of Spillover and Spillback in SARS-CoV-2 Transmission and the Importance of One Health in Understanding the Dynamics of the COVID-19 Pandemic. Journal of Clinical Microbiology, e01610–22. https://doi.org/10.1128/jcm.01610-22

Vandegrift, K. J., Yon, M., Surendran Nair, M., Gontu, A., Ramasamy, S., Amirthalingam, S., Neerukonda, S., Nissly, R. H., Chothe, S. K., Jakka, P., LaBella, L., Levine, N., Rodriguez, S., Chen, C., Sheersh Boorla, V., Stuber, T., Boulanger, J. R., Kotschwar, N., Aucoin, S. G., … Kuchipudi, S. V. (2022). SARS-CoV-2 Omicron (B.1.1.529) Infection of Wild White-Tailed Deer in New York City. Viruses, 14(12), 2770. https://doi.org/10.3390/v14122770

Willgert, K., Didelot, X., Surendran-Nair, M., Kuchipudi, S. V., Ruden, R. M., Yon, M., Nissly, R. H., Vandegrift, K. J., Nelli, R. K., Li, L., Jayarao, B. M., Levine, N., Olsen, R. J., Davis, J. J., Musser, J. M., Hudson, P. J., Kapur, V., & Conlan, A. J. K. (2022). Transmission history of SARS-CoV-2 in humans and white-tailed deer. Scientific Reports, 12(1), 12094. https://doi.org/10.1038/s41598-022-16071-z

Zhao, J., Cui, W., & Tian, B. (2020). The Potential Intermediate Hosts for SARS-CoV-2. Frontiers in Microbiology, 11, 580137. https://doi.org/10.3389/fmicb.2020.580137

Zhou, H., Chen, X., Hu, T., Li, J., Song, H., Liu, Y., Wang, P., Liu, D., Yang, J., Holmes, E. C., Hughes, A. C., Bi, Y., & Shi, W. (2020). A Novel Bat Coronavirus Closely Related to SARS-CoV-2 Contains Natural Insertions at the S1/S2 Cleavage Site of the Spike Protein. Current Biology, 30(11), 2196–2203.e3. https://doi.org/10.1016/j.cub.2020.05.023

